# Airway smooth muscle tone curbs hyperresponsiveness in experimental asthma

**DOI:** 10.1101/2024.07.05.602208

**Authors:** Morgan Gazzola, Magali Boucher, Cyndi Henry, Andrés Rojas-Ruiz, David Marsolais, Ynuk Bossé

## Abstract

**Background & Objectives:** A sustained contraction of airway smooth muscle, hereinafter called tone, increases the response to methacholine in healthy mice and humans. However, the effect of tone in the context of an active inflammation remains to be investigated. The objective of the present study was to test the effect of tone on the *in vivo* response to methacholine in mice during an active inflammatory phase of experimental asthma.

**Methods:** Male BALB/c mice were exposed once-daily to either intranasal saline or house dust mite for 10 consecutive days to induce experimental asthma. They then underwent one of two methacholine challenges 24 h after the last exposure. While the same cumulative dose was administered in both challenges, one was preceded by a 20-min period of tone induced by nebulizing low doses of methacholine at 5-min intervals. Respiratory mechanics were monitored before and throughout the methacholine challenge by oscillometry. Bronchoalveolar lavages (BAL) and histology were also performed.

**Results:** BAL inflammation and histological alterations were consistent with experimental asthma. In accordance with previous studies, tone potentiated the response to methacholine in control mice, mainly by stiffening the lung periphery. The lung was even stiffer upon methacholine challenge during an active phase of inflammation in mice with experimental asthma, but this was not further potentiated by tone. In fact, in mice with experimental asthma, tone mitigated hyperresponsiveness by preventing further airway narrowing and, more importantly, small airway narrowing heterogeneity and closure.

**Conclusion:** During an active inflammatory phase of experimental asthma, tone protects against hyperresponsiveness.

**Take-home message:** The effect of airway smooth muscle tone on the methacholine response was investigated in mice with or without experimental asthma. While tone potentiated the methacholine response in control mice, it mitigated hyperresponsiveness in experimental asthma. These results unveiled a protective role of the airway smooth muscle in experimental asthma.

## INTRODUCTION

Tone, defined herein as a sustained contraction, increases the contractile capacity of airway smooth muscle (ASM) *in vitro* through a process dubbed ‘force adaptation’^1^. The molecular mechanisms governing force adaptation are not fully delineated but involve actin filamentogenesis^2^. Force adaptation occurs in sheep tracheal ASM strips^1, 3, 4^, tracheas from C57BL/6 mice^5^, human bronchi^6^, and in isolated human ASM cells^2^. Recent experiments also demonstrated that force adaptation occurs to the same extent in ASM derived from both sexes in BALB/c mice^7^. Force adaptation is thus a universal phenomenon occurring in all tested species and in both males and females.

Investigating force adaptation *in vivo* is not as straightforward. Yet, four studies have previously investigated the effect of tone on the *in vivo* response to methacholine^5–8^. In young healthy humans and mice, tone potentiates the response to methacholine^5–8^. In-depth analyses of respiratory mechanics further suggested that, in both mice and humans, the potentiating effect of tone on the methacholine response is substantially greater on small peripheral airways and lung stiffness compared to large airways^6, 7^. These findings are relevant to asthma, which is characterized by an elevated tone^9, 10^ and hyperresponsiveness to methacholine largely due to small airway dysfunction^11–16^. However, further analyses in mice demonstrated that the stiffening of the lung caused by tone protects against small airway narrowing heterogeneity^7^. These results alternatively suggested that tone may potentially be beneficial in lung disorders displaying a propensity for airway narrowing heterogeneity, such as asthma^15, 17–20^. The goal of the present study is to investigate the effect of tone on the *in vivo* response to methacholine in a mouse model during an inflammatory phase of experimental asthma.

## METHODS

### Mice

Male BALB/c mice (Charles River, Saint-Constant, Canada) were tested at 8 weeks of age. Males were chosen because the effect of ASM contraction on their lung mechanics is substantially greater than females^7, 21, 22^. They were provided food and water *ad libitum*. All procedures were approved by the Committee of Animal Care of *Université Laval* in accordance with guidelines of the Canadian Council on Animal Care (protocols 2018-005-4 and 2022-977-1).

### Experimental asthma

Mice were divided into two groups of 44 (Figure 1A): one group exposed to 25 μL of saline (control mice) and one group exposed to 25 μL of 2 mg/mL of house dust mite (HDM) extract (*D. pteronyssinus*; Greer, Lenoir, USA) to induce pulmonary allergic inflammation (inflamed mice). The endotoxin concentration was 47.3 EU per mg of HDM extract. They were exposed through an intranasal instillation once daily under isoflurane anesthesia for 10 consecutive days. All outcomes were measured 24 h after the last exposure.

**Figure 1.**
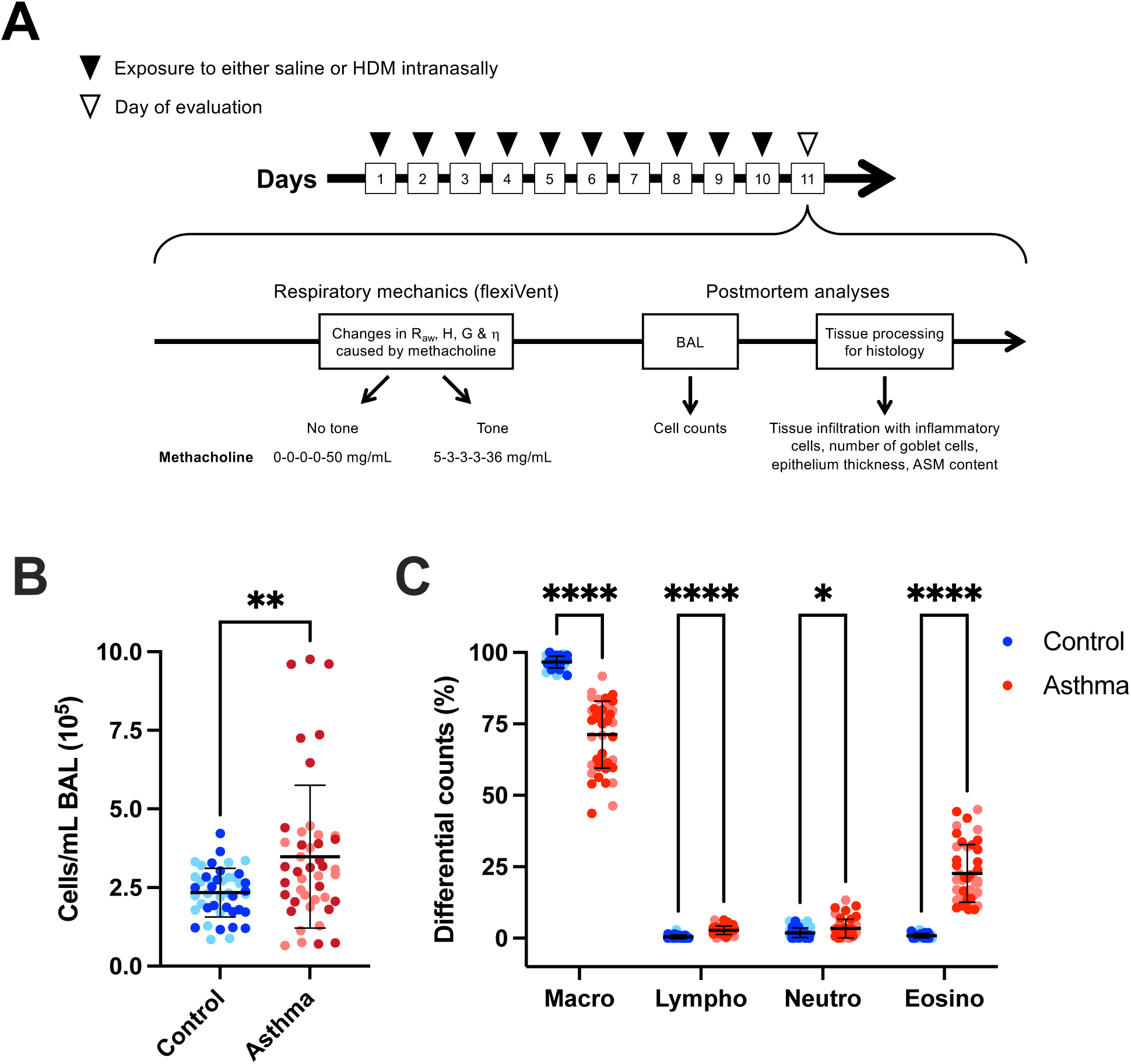
**(A)** Schematic depicting the sequence of interventions in the mouse study. See Methods for further details. **(B)** The number of total inflammatory cells per mL of bronchoalveolar lavages is shown for control (blue) and inflamed (red) mice. **(C)** The differential cell counts in bronchoalveolar lavages expressed in percentages are shown for control (blue) and inflamed (red) mice. Data are individual results, together with the mean ± SD for each group. The faded and bright colors in each group refer to mice exposed to the methacholine challenge without and with tone, respectively. Significant differences are indicated by asterisks (*, ** and **** are p<0.05, 0.01 and 0.0001, respectively). N = 44 (22 exposed to tone and 22 not exposed to tone) Abbreviations: ASM, airway smooth muscle; BAL, bronchoalveolar lavages; Eosino, eosinophils; ρι (eta), hysteresivity; G, tissue resistance; H, lung elastance; HDM, house dust mite; Lympho, lymphocytes; Macro, macrophages; Neutro, neutrophils; and R_aw_, airway resistance

### Respiratory mechanics

Mice were prepared and ventilated with the flexiVent (FX Module 2, SCIREQ, Montreal, Canada) as previously described^23–26^. Briefly, they were anesthetized and put under systemic analgesia using ketamine (100 mg/kg) and xylazine (10 mg/kg). They were then tracheotomized and connected to the flexiVent in a supine position. They were ventilated mechanically at a tidal volume of 10 mL/kg with an inspiratory-to-expiratory time ratio of 2:3 at a breathing frequency of 150 breaths/min and with a positive end expiratory pressure of 3 cmH_2_O. To avoid spontaneous breathing, they were also paralyzed by injecting 100 and 300 µL of pancuronium bromide (0.1 mg/kg) intramuscularly and intraperitoneally, respectively.

Respiratory mechanics were also assessed with the flexiVent, using an oscillometric perturbation called the Quick Prime-3. This perturbation was actuated twice at baseline and 7 times at 38-s intervals starting 30 s after the end of each bout of nebulization. The only exception was after the final dose of methacholine, where it was actuated 15 times at 20-s intervals, starting again 30 s after the end of nebulization. The volume perturbation imposed by the Quick Prime-3 is a composite input flow signal, made of 13 sine waves of mutually prime frequencies with different amplitudes and phases, which enables the calculation of impedance of the respiratory system based on the resulting output pressure signal^27^. Impedance was then analyzed using a computational model called the constant phase model to deduce three parameters^28^. One is airway resistance (R_aw_), representing the resistance to airflow in conducting airways. Its change in response to methacholine (ΔR_aw_) is a surrogate for airway narrowing. Another one is lung elastance (H), representing the elastance of the lung and the chest wall. Its change in response to methacholine (ΔH) is mainly due to both lung tissue stiffening and small airway closure. The other one is tissue resistance (G), representing the resistance of the lung and the chest wall. Similar to ΔH, its change in response to methacholine (ΔG) is sensitive to lung tissue stiffening and small airway closure. However, and unlike ΔH, it is also very sensitive to small airway narrowing heterogeneity^29^. The change in hysteresivity (ρι), which is the ratio of G over H, in response to methacholine (Δη) is thus an index of airway narrowing heterogeneity^30^.

### Methacholine challenges

For each bout of nebulization, the nebulizer for small particle size (Aeroneb Lab, Aerogen Inc, Galway, Ireland) was operating for a duration of 10 s at a duty cycle of 50% under regular ventilation. The effect of tone was tested as previously described^5, 7^, except that doses were reduced to accommodate the greater response to methacholine in inflamed mice. Briefly, for both the control and inflamed groups, the response to methacholine was compared between two subgroups of mice subjected to a nebulized challenge that started with a 20-min period either without or with tone (Figure 1A). Tone was maintained for 20 min by administering 4 low doses of methacholine at 5-min intervals. Doses were 5, 3, 3 and 3 mg/mL. The subgroup without tone received nebulized saline 4 times at 5-min intervals. After 20 min, a high dose of methacholine was then nebulized. Importantly, this final high dose was adjusted so that each mouse, exposed or not to tone, received the same cumulative dose of methacholine. The final dose was either 36 or 50 mg/mL in subgroups exposed or not to tone, respectively, for a total cumulative dose of 50 mg/mL. In all mice, the methacholine response was assessed by measuring the changes in oscillometric readouts (H, G, ρι & R_aw_) from their baseline values (*i.e.*, at the beginning of the challenge at time zero).

### Bronchoalveolar lavages (BAL)

One mL of phosphate-buffered saline (PBS) was injected in the lung through the trachea and aspirated to recover the BAL. This was repeated 3 times and the recovered BAL were pooled together. The total volume was recorded and centrifuged at 500-x g for 5 min. The supernatant was discarded and the pellet was resuspended in 200 μL of PBS for control mice and 500 μL for inflamed mice. Total cells in BAL were stained with crystal violet and counted using a hemacytometer. Seventy-five thousand cells were also cytospun and stained with modified May-Grünwald Giemsa to count macrophages, lymphocytes, neutrophils and eosinophils.

### Histology

Histology was performed as previously described^26, 31^ on the left lung of 10 mice in each group. Briefly, the lung was immersed in formalin during 24 h for fixation. The formalin was replaced by progressively upraising the ethanol concentration to dehydrate the tissue. The lung was then embedded in paraffin and cut in 5 μm-thick sections that were subsequently deposited on microscopic slides and stained with hematoxylin and eosin (H&E), Periodic acid-Schiff (PAS) with alcian blue, or Masson trichrome. The slides were then scanned with a NanoZoomer Digital scanner (Hamamastu photonics, Bridgewater, USA) at 40X.

H&E stain was performed to evaluate the infiltration of inflammatory cells within the lung tissue. Fifteen non-overlapping photomicrographs (1440 × 904 pixels) from 3 non-contiguous lung sections were blindly scored from zero (no inflammation) to 5 (very severe inflammation).

PAS with alcian blue was used to count the number of goblet cells. All airways cut transversally in 3 non-contiguous lung sections were analyzed, representing 5 to 16 airways per mouse (average of 10.8 ± 2.6). The number of goblet cells within each airway was divided by the length of the basement membrane. The same staining was used to measure the epithelium thickness. All airways cut transversally and displaying a full circumference in 3 non-contiguous lung sections were analyzed, representing 3 to 16 airways per mouse (average of 10.5 ± 2.7). The thickness was analyzed by measuring the area occupied by the epithelium divided by the basement membrane perimeter.

Masson trichrome was used to quantify the content of smooth muscle within the airway wall. All airways cut transversally in 3 non-contiguous lung sections were analyzed, representing 2 to 14 airways per mouse (average of 8.7 ± 2.8). The content of ASM in each airway was calculated by measuring the area occupied by ASM divided by the square of its basement membrane perimeter.

### Data analysis

Since data were often not normally distributed, nonparametric analyses were performed. Two-factor ART ANOVAs were used to assess the effect of inflammation, tone, and their interaction on the response to methacholine. When the interaction was significant, it was followed by Mann-Whitney tests corrected for multiple comparisons using the Holm-Sidak method to specifically assess the effect of tone on the methacholine response in control and inflamed mice separately. Mann-Whitney tests were also used to compare the effect of tone (at the 20-min time point, just before adding the final dose) between control and inflamed mice. Since tone had no effect on BAL cells and histology, mice exposed or not to tone were combined and the different readouts were compared between control and inflamed mice using Mann-Whitney tests. All statistical analyses were performed using Prism 10 (version 10.2.3, GraphPad, San Diego, USA), except two-factor ART ANOVAs, which were done in R. p<0.05 was considered significant.

## RESULTS

The number of total cells/mL of bronchoalveolar lavages is depicted in Figure 1B. Experimental asthma increased the number of total cells/mL from 2.3 ± 0.8 to 3.5 ± 2.3 × 10^5^ (p=0.007). Differential cell counts expressed in percentage of total cells are depicted in Figure 1C. Experimental asthma significantly decreased the percentage of macrophages from 96.7 ± 2.1 to 71.3 ± 11.7% (p<0.0001), and significantly increased the percentage of: 1-lymphocytes from 0.5 ± 0.6 to 2.8 ± 1.5% (p<0.0001); 2-neutrophils from 1.9 ± 1.7 to 3.4 ± 3.2% (p=0.012); and 3-eosinophils from 0.9 ± 0.8 to 22.7 ± 10.1% (p<0.0001).

The infiltration of the lung tissue with inflammatory cells in control and inflamed mice is depicted in Figure 2A. Experimental asthma increased cellular infiltration from a score of 2.1 ± 0.3 to 3.3 ±0.3 (p<0.0001). Goblet cell counts and the epithelium thickness are depicted in Figure 2B. Experimental asthma increased the count of goblet cells/mL of basement membrane from 1.1 ±0.8 to 66.4 ± 24.2 (p<0.0001) and the epithelium thickness from 17.4 ± 1.7 to 24.5 ± 3.1 µm (p<0.0001). The content of ASM within the airway wall is depicted in Figure 2C and was not affected by experimental asthma.

**Figure 2.**
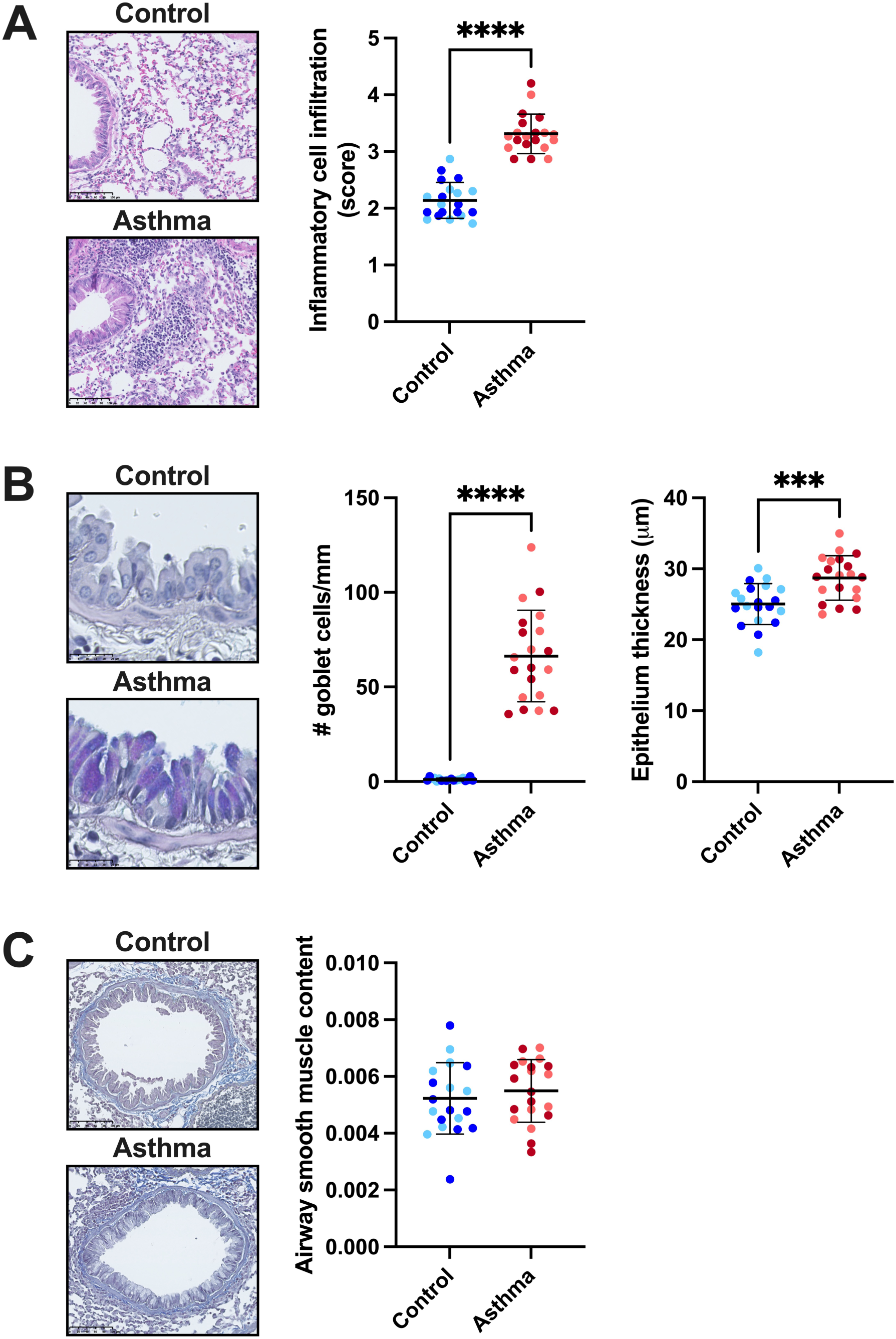
Lung histology. Images on the left are representative images of lung sections. Results from control (blue) and asthmatic (red) mice are shown in scatter plots on the right. **(A)** Staining with H&E showing the infiltration of the lung tissue with inflammatory cells. Scale bar is 100 µm. For each image, an arbitrary inflammatory score ranging from 0 (no cellular infiltration) to 5 (very severe cellular infiltration) was assigned. **(B)** Staining with PAS and alcian blue showing goblet cells and the epithelium thickness. Scale bar is 25 µm. For each airway, the number of goblet cells/basement membrane length and the area occupied by the epithelium/basement membrane perimeter (*i.e.*, epithelium thickness) were calculated. **(C)** Staining with Masson trichrome showing the content of smooth muscle within the airway wall. Scale bar is 100 µm. For each bronchus, the area occupied by airway smooth muscle was divided by the square of the basement membrane perimeter. Data in scatter plots are individual results, together with the mean ± SD for each group. The faded and bright colors in each group refer to mice exposed to the methacholine challenge without and with tone, respectively. Significant differences are indicated by asterisks (*** and **** is p<0.001 and 0.0001, respectively). N = 20 (10 not exposed to tone and 10 exposed to tone).

The changes in lung elastance (ΔH) over the entire methacholine challenges are depicted in Figure 3A (left and right panels for control and inflamed mice, respectively). After 20 min, prior to the administration of the final dose, tone increased H by 12.1 ± 5.7 cmH_2_O·s/mL in control mice and by 26.0 ± 22.9 cmH_2_O·s/mL in inflamed mice. The difference between control and inflamed mice was significant (p=0.031), confirming hyperresponsiveness. The AUC over 5 min after the introduction of the final dose of methacholine is depicted in Figure 3B. The AUC-ΔH was increased by inflammation (p<0.0001), and there was also a significant interaction between inflammation and tone (p=0.011) (Figure 3B). Post-hoc analyses show that tone increased the AUC-ΔH but only in control mice (p<0.0001).

**Figure 3.**
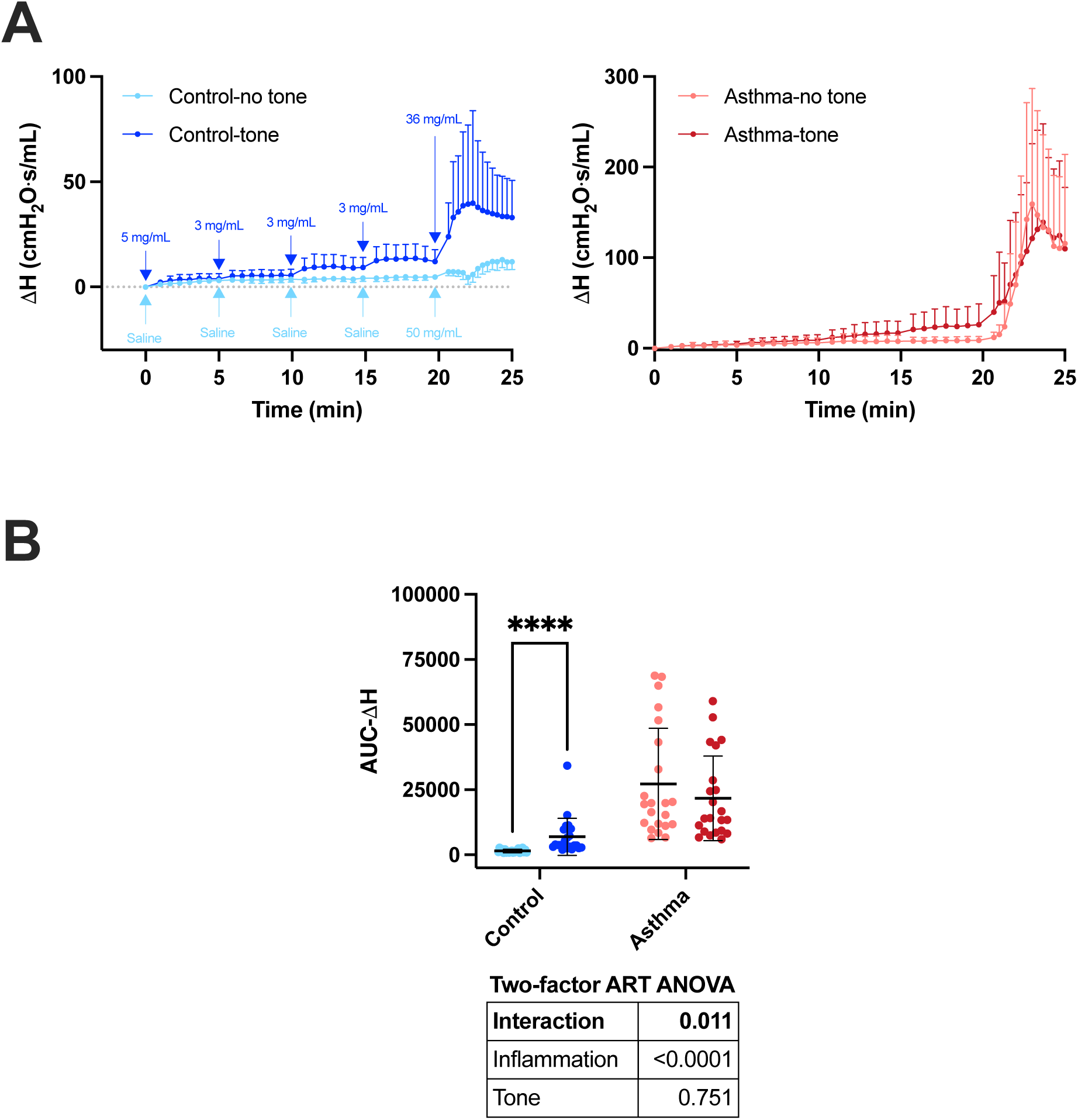
Tone on the methacholine response in terms of changes in lung elastance (H). **(A)** Methacholine-induced changes in H from baseline (ΔH) in control (blue on the left panel) and inflamed mice (red on the right panel) are shown for the entire challenges, including the 20 min period without (faded colors) or with (bright colors) tone. The dosing regimen is displayed in control mice. The same regimen was used for inflamed mice. Note that the scale for the y-axis is markedly different between control and inflamed mice, clearly showing that mice with experimental asthma were hyperresponsive to methacholine. **(B)** The methacholine response in each mouse was calculated by measuring the area under the curve (AUC) over 5 min after the administration of the final dose (AUC-ΔH). Individual results, together with the mean ± SD, are shown for each group. Results of a two-factor ART ANOVA are shown in the table below the graph. Significant differences in post-hoc tests are indicated by asterisks (**** is p<0.0001). N = 22

The changes in tissue resistance (ΔG) over the entire methacholine challenges are depicted in Figure 4A. After 20 min, prior to the administration of the final dose, tone increased G by 2.73 ±1.12 cmH_2_O·s/mL in control mice and by 6.46 ± 6.47 cmH_2_O·s/mL in inflamed mice. The difference between control and inflamed mice was not significant (p=0.058). The AUC over 5 min after the introduction of the final dose of methacholine is depicted in Figure 4B. The AUC-ΔG was increased by inflammation (p<0.0001) and tone (p=0.024), and there was also a significant interaction between inflammation and tone (p=0.0009) (Figure 4B). Post-hoc analyses show that tone increased the AUC-ΔG but only in control mice (p=0.023).

**Figure 4.**
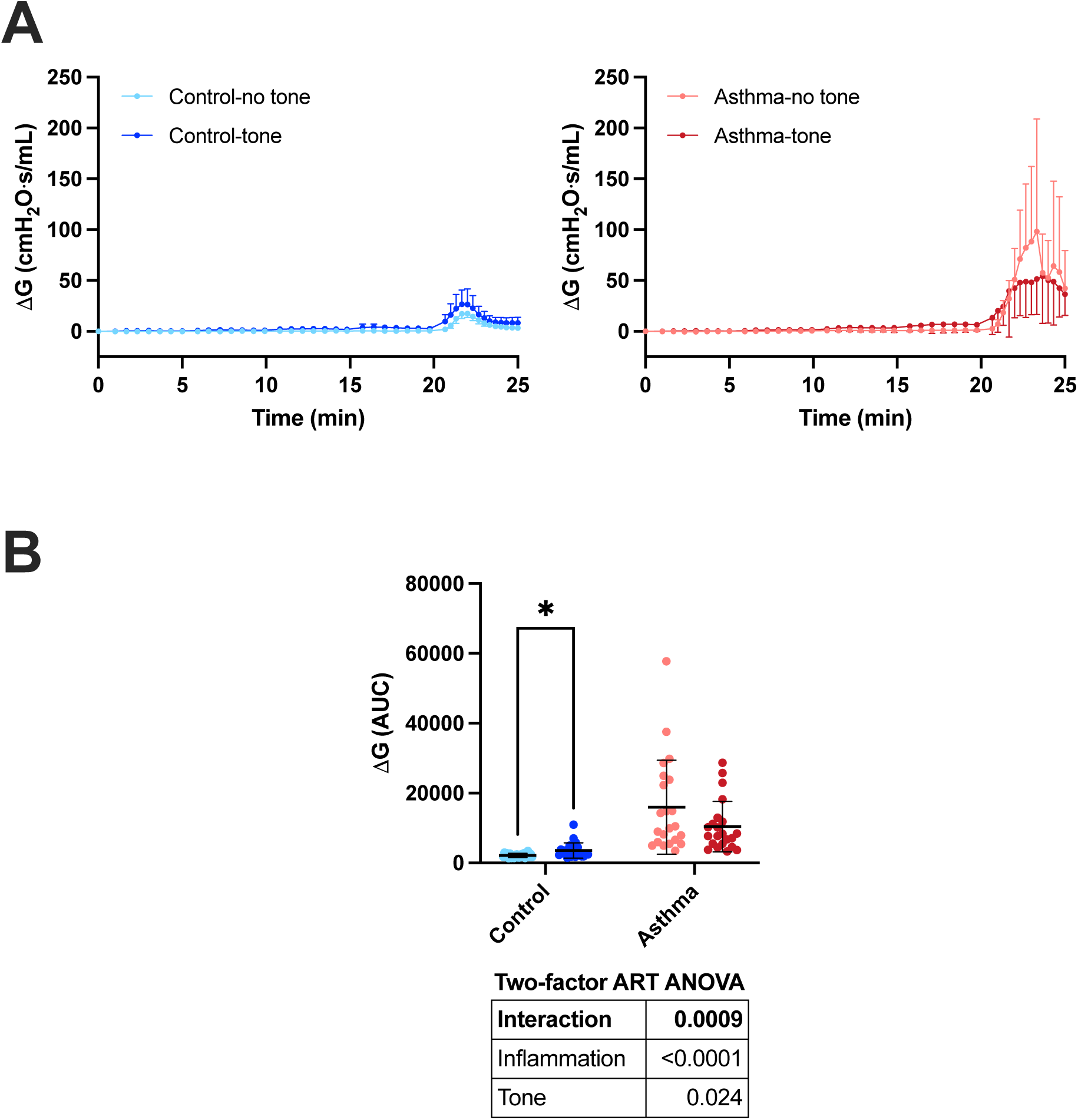
Tone on the methacholine response in terms of changes in tissue resistance (G). **(A)** Methacholine-induced changes in G from baseline (ΛG) in control (blue on the left panel) and inflamed mice (red on the right panel) are shown for the entire challenges, including the 20 min period without (faded colors) or with (bright colors) tone. The dosing regimen is depicted Figure 3. **(B)** The methacholine response in each mouse was calculated by measuring the area under the curve (AUC) over 5 min after the administration of the final dose (AUC-ΛG). Individual results, together with the mean ± SD, are shown for each group. Results of a two-factor ART ANOVA are shown in the table below the graph. Significant differences in post-hoc tests are indicated by asterisks (* is p<0.05). N = 22

Airway narrowing heterogeneity over the entire methacholine challenges, assessed by measuring the changes in hysteresivity (Δη), is depicted in Figure 5A. After 20 min, prior to the administration of the final dose, tone slightly increased ρι by 0.015 ± 0.015 in control mice and by 0.029 ± 0.037 in inflamed mice. The difference between control and inflamed mice was not significant (p=0.493). The AUC over 5 min after the introduction of the final dose of methacholine is depicted in Figure 5B. The AUC-Δη, used as an index of airway narrowing heterogeneity, was increased by inflammation (p<0.0001) and decreased by tone (p<0.0001), and there was also a significant interaction between inflammation and tone (p=0.046) (Figure 5B). Post-hoc analyses show that tone decreased AUC-Δη in both control (p=0.009) and inflamed mice (p=0.006).

**Figure 5.**
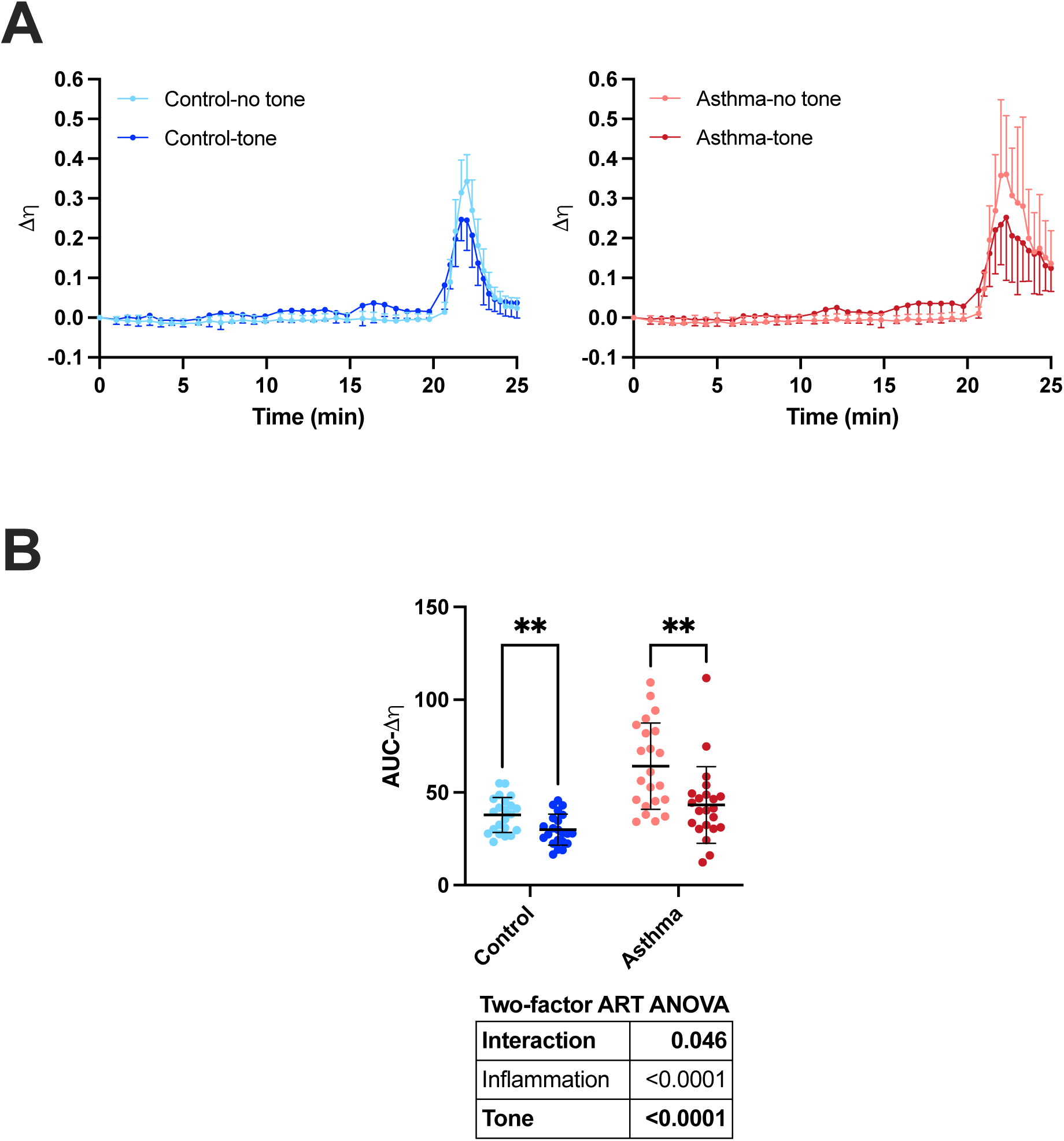
Tone on the methacholine response in terms of changes in hysteresivity (ρι). **(A)** Methacholine-induced changes in ρι from baseline (Δη) in control (blue on the left panel) and inflamed mice (red on the right panel) are shown for the entire challenges, including the 20 min period without (faded colors) or with (bright colors) tone. The dosing regimen is depicted in Figure 3. **(B)** The methacholine response in each mouse was calculated by measuring the area under the curve (AUC) over 5 min after the administration of the final dose (AUC-Δη). Individual results, together with the mean ± SD, are shown for each group. Results of a two-factor ART ANOVA are shown in the table below the graph. Significant differences in post-hoc tests are indicated by asterisks (** is p<0.01). N = 22

Airway narrowing over the entire methacholine challenges, assessed by measuring the changes in airway resistance (ΔR_aw_), is depicted in Figure 6A. After 20 min, prior to the administration of the final dose, tone increased R_aw_ by 0.14 ± 0.09 cmH_2_O·s/mL in control mice and by 0.20 ± 0.16 cmH_2_O·s/mL in inflamed mice. The difference between control and inflamed mice was not significant (p=0.634). The AUC over 5 min after the introduction of the final dose of methacholine is depicted in Figure 6B. The AUC-ΔR_aw_, used as an index of airway narrowing, was increased by inflammation (p=0.0003) and also significantly affected by tone (p=0.007). There was also a significant interaction between inflammation and tone (p=0.003) (Figure 6B). Post-hoc analyses show that tone decreased AUC-ΔR_aw_ but only in inflamed mice (p=0.042). More precisely, tone reduced hyperresponsiveness (*i.e.*, the difference in the methacholine response between control and inflamed mice) by a mean of 86.3% (Figure 6B).

**Figure 6.**
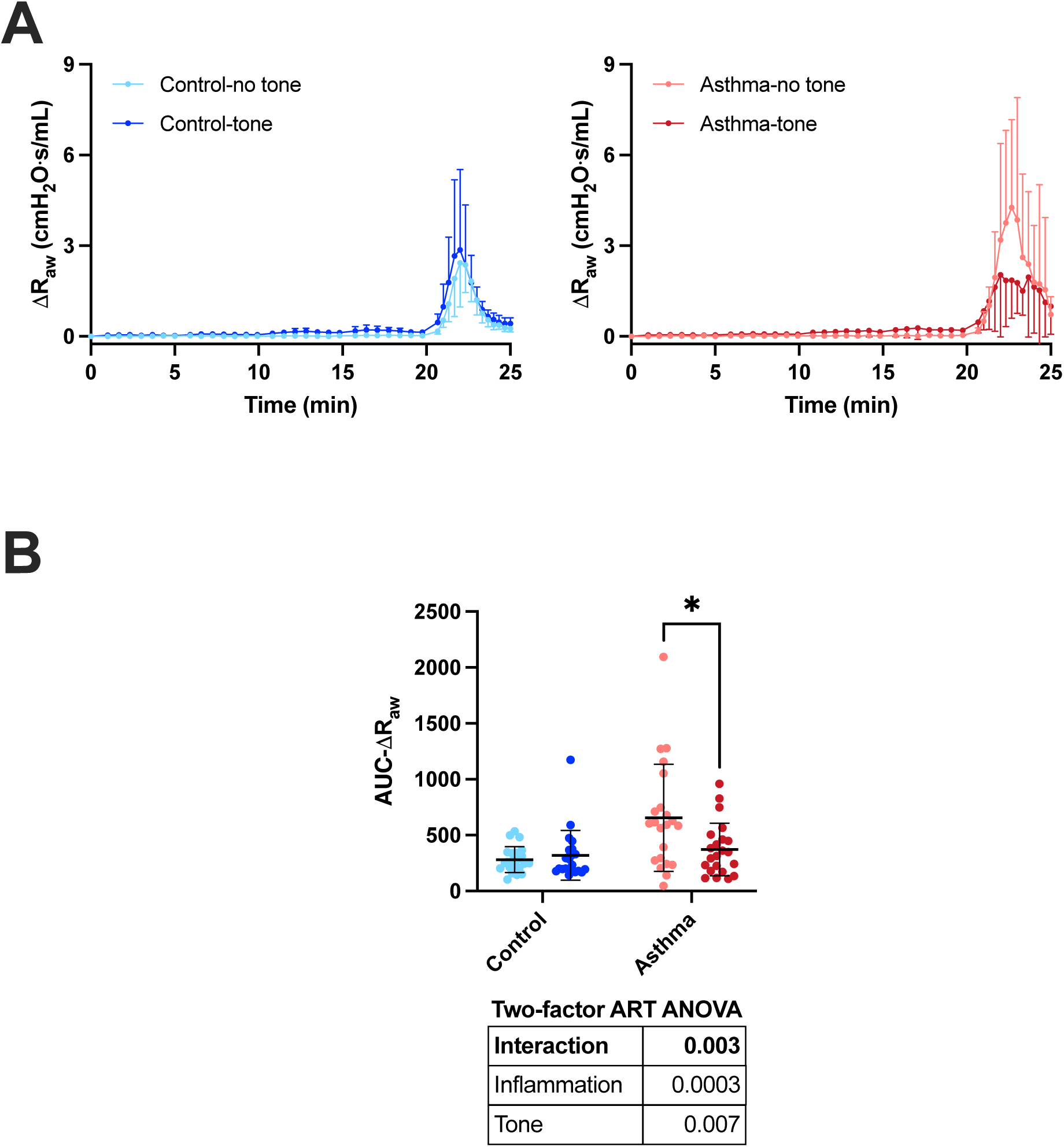
Tone on the methacholine response in terms of changes in airway resistance (R_aw_). **(A)** Methacholine-induced changes in R_aw_ from baseline (ΔR_aw_) in control (blue on the left panel) and inflamed mice (red on the right panel) are shown for the entire challenges, including the 20 min period without (faded colors) or with (bright colors) tone. The dosing regimen is depicted in Figure 3. **(B)** The methacholine response in each mouse was calculated by measuring the area under the curve (AUC) over 5 min after the administration of the final dose (AUC-Δ R_aw_). Individual results, together with the mean ± SD, are shown for each group. Results of a two-factor ART ANOVA are shown in the table below the graph. Significant differences in post-hoc tests are indicated by asterisks (** is p<0.05). N = 22

## DISCUSSION

This study demonstrated that tone protects against methacholine hyperresponsiveness in mice with experimental asthma. It represents the first *in vivo* evidence of a protective effect of ASM contraction in the context of an active inflammation. It is a discovery that is changing our perception on the role played by ASM in respiratory mechanics, especially in diseases.

Several studies have demonstrated that tone increases the contractile capacity of ASM^1, 3–7, 32^. How this phenomenon, called force adaptation, integrates into the whole respiratory system to influence the *in vivo* response to methacholine was difficult to predict^33, 34^. The present study is consistent with previous studies on healthy subjects showing that tone potentiates the methacholine response^5–8^. However, when the effect of tone on the methacholine response was compared in mice with and without experimental asthma, a significant interaction was observed between inflammation and tone. This was true irrespective of the readouts used to monitor the methacholine response (H, G, ρι, or R_aw_). It indicates that the effect of tone on the response to methacholine depended on whether it was tested in control or inflamed mice. More precisely, while tone increased the response to methacholine in control mice, it rather attenuated the hyperresponsiveness to methacholine in inflamed mice.

The fine-grained physiological analyses undertaken herein allowed us to speculate about the underlying mechanisms. Note that tone and inflammation both potentiated the methacholine response of H when acting alone, but then exerted no additive or synergic effect when acting together (Figure 3B). The methacholine response of H in inflamed mice exposed to tone thus seems to rely on two competing mechanisms, namely a stiffening of the lung that prevented small airway closure. Indeed, H is sensitive to both lung stiffening and airway closure^30^, the latter being one of the main causes of hyperresponsiveness in mouse models of asthma^35^. Therefore, while the stiffening of the lung caused by tone may have increased H in control mice, it may have concurrently mitigated small airway closure in inflamed mice, thereby explaining the lack of effect of tone on the methacholine response of H in inflamed mice (Figure 3B). Concordantly, other studies have shown that methacholine stiffens the lung tissue^25, 36^. Our previous study in control mice has also shown that the potentiating effect of tone on the methacholine response of H was due to a real stiffening of the lung tissue and not to airway closure^7^.

The functional consequence of lung stiffening caused by tone was also confirmed otherwise. First, tone prevented airway narrowing heterogeneity. This was shown in both control and inflamed mice (Figure 5B). It is important to understand that heterogeneity is a precursor of closure^29, 37^; *i.e.*, greater heterogeneity invariably leads to greater closure. Airway narrowing is also inherently heterogeneous^20, 38–44^. Therefore, it would be very unlikely that tone had decreased heterogeneity without reducing closure. As per our previous study^7^, the protective effect of tone on airway narrowing heterogeneity was shown by measuring the methacholine-induced changes in hysteresivity (Figure 5). Hysteresivity is the ratio of G over H and is a readout indisputably sensitive to narrowing heterogeneity^30^. This is because while both G and H are sensitive to closure, G is also sensitive to airway narrowing heterogeneity^29^. Therefore, while airway closure increases G and H in the same proportion, leaving hysteresivity intact, narrowing heterogeneity further increases G and, thereby, increases hysteresivity. In our study, the protective effect of tone on airway narrowing heterogeneity was even stronger in inflamed *versus* control mice (interaction; p=0.046), presumably due to an increased propensity for heterogeneity in diseases^17–20, 29, 41, 45^.

Second, tone decreased methacholine-induced airway narrowing in inflamed mice (Figure 3D). This effect was not small. Tone reduced by a mean of 86% the potentiating effect of inflammation on the narrowing induced by methacholine. This ultimately proved that tone protects against hyperresponsiveness in the context of inflammation. We think that this was mainly due to lung stiffening as well, since stiffening should prevent narrowing by increasing the load impeding ASM shortening.

The reason this function of ASM has been missed out that many often is probably a question of timing (kinetics). In the present study, serial low doses of methacholine were nebulized at 5-min intervals during 20 min. This probably allowed sufficient time for the methacholine to dissolve across the lung. It is probably when ASM throughout the whole lung is activated, especially the one in small airways, that a homogenous lung stiffening ensues and protects against further airway narrowing, heterogeneity and closure.

Our results are consistent with others. Brown and colleagues have demonstrated in dogs that challenging the whole lung with histamine is protective against airway narrowing compared to delivering histamine locally to a few specified airways^46^. Again, this is attributed to the force of interdependence between airways; *i.e.*, narrowing of neighboring airways promote dilation. Studies testing bronchodilator drugs *in vivo* in humans also supported the role of ASM in preserving small airway patency. Crawford and coworkers, for example, found that relaxing ASM *in vivo* in healthy subjects increases ventilation heterogeneity^47^. The study of Kelly and coworkers substantiated this finding by demonstrating that relaxing ASM in healthy subjects increases residual volume (RV) and RV/total lung capacity (TLC)^10^, the latter indicating air trapping due to closed airways. Notably, oscillometric readouts reflecting airway caliber, such as resistance at 5 Hz, were improved by the bronchodilator drug in that latter study^10^. These results suggest that, as contrasting as that may sound, a bronchodilator drug can dilate large airways while concurrently closing small airways (consistent with the notion that constriction protects against small airway narrowing and closure). Combined with our study, these results suggest that while an incipient contraction of ASM almost inevitably reduces airway caliber, a small level of constriction in all airways, together with the ensuing stiffening of the lung, prevents further airway narrowing, heterogeneity and closure.

The effect of tone in control and inflamed mice thus seem contrasting because it acts on different determinants of the methacholine response. In healthy subjects, the effect of tone seems detrimental because the gain in ASM’s contractile capacity through force adaptation^1, 3–7, 32^ translated into a slight increase in airway narrowing combined with a considerable stiffening of the lung^6, 7^. However, during inflammation, the effect of tone is beneficial because it prevented excessive airway narrowing and, most importantly, small airway narrowing heterogeneity and closure, the latter being the two main determinants of asthmatic hyperresponsiveness^17–20, 29, 35, 41, 45, 48, 49^. We surmise that by preventing small airway closure, a homogeneous stiffening of the lung by ASM tone may represent a good compromise to preserve ventilation in inflamed pulmonary diseases.

## Conclusion

The results of the present study suggest that the seemingly detrimental effects of tone seen in control mice is outweighed in inflamed mice by its protecting effect on airway narrowing, as well as on small airway narrowing heterogeneity and closure. Together, these results suggest that

ASM tone may be protective in asthma, urging us to revisit one of the therapeutic goals in asthma that aims at relaxing the ASM.

## DATA AVAILABILITY STATEMENT

The datasets used and analyzed during the current study are available from the corresponding author on reasonable request.

